# Spike frequency adaptation in primate lateral prefrontal cortex neurons results from interplay between intrinsic properties and circuit dynamics

**DOI:** 10.1101/2024.09.03.610998

**Authors:** Nils A. Koch, Benjamin W. Corrigan, Michael Feyerabend, Roberto A. Gulli, Michelle S. Jimenez-Sosa, Mohamad Abbass, Julia K. Sunstrum, Sara Matovic, Megan Roussy, Rogelio Luna, Samuel A. Mestern, Borna Mahmoudian, Susheel Vijayraghavan, Hiroyuki Igarashi, Kartik S. Pradeepan, William J. Assis, J. Andrew Pruszynski, Shreejoy Tripathy, Jochen F. Staiger, Guillermo Gonzalez-Burgos, Andreas Neef, Stefan Treue, Stefan Everling, Wataru Inoue, Anmar Khadra, Julio C. Martinez-Trujillo

## Abstract

Recordings of cortical neurons isolated from brain slices and dissociated from their networks, display intrinsic spike frequency adaptation (I-SFA) to a constant current input. Interestingly, extracellular recordings in behaving subjects also show extrinsic-SFA (E-SFA) in response to sustained visual stimulation. Because neurons are isolated from brain networks in slice recordings, it is challenging to infer how I-SFA contributes to E-SFA in interconnected brains during behavior. To investigate this, we recorded responses of macaque lateral prefrontal cortex neurons *in vivo* during a visually guided saccade task and in acute brain slices *in vitro*. Broad spiking (BS) putative pyramidal cells and narrow spiking (NS) putative inhibitory interneurons exhibited E-SFA *in vivo*. In acute brain slices, both cell types displayed I-SFA though their magnitudes differed. To investigate how *in vitro* I-SFA contributes to *in vivo* E-SFA, we developed a data-driven hybrid circuit model in which local NS neurons are driven by BS input. We observed that model NS cell responses show longer SFA than observed *in vivo*. Introducing inhibition of NS cells to the model circuit removed this discrepancy. Our results indicate that both I-SFA and inhibitory circuit dynamics contribute to E-SFA in LPFC neurons. They highlight the contribution of single neuron and network dependent computations to neural activity underlying behavior.

## Introduction

Previous studies have shown that single neurons in the neocortex are not linear filters but they exhibit time varying firing patterns in response to constant stimulation depending on the cell’s intrinsic membrane properties (Ascoli et al., 2008). One contributor to such diversity of patterns is intrinsic spike frequency adaptation (I-SFA), the temporal decay in the firing rate of a neuron in response to a constant stimulus (Benda, 2021; Ha & Cheong, 2017). In insect and rodent sensory systems, I-SFA serves as a high-pass filter (Benda, 2021; Benda et al., 2005; Prescott & Sejnowski, 2008) that produces input intensity invariance (Benda & Hennig, 2008; Clemens et al., 2018; Kastner & Baccus, 2014; Peron & Gabbiani, 2009), encodes rate-of-change (Lee et al., 2023), allows for stimulus specific adaptation (Benda, 2021; Whitmire & Stanley, 2016), and generates sparse neural codes (Betkiewicz et al., 2020). In primates, SFA has been documented in sensory cortical areas such as visual area MT using extracellular recordings in behaving and anesthetized monkeys (Lisberger & Movshon, 1999; Priebe et al., 2002). We will refer to this form of adaptation as extrinsic-SFA (E-SFA). Although, E-SFA has been less studied in executive control areas of primates, a few studies have described phasic or adapting responses to visual stimulus onsets in the lateral prefrontal cortex (LPFC) (Bullock et al., 2017; Funahashi et al., 1990; Mikami et al., 1982). The latter suggests that E-SFA is a ubiquitous feature of neuronal responses across the neocortex. The factors that contribute to E-SFA across different neuronal types in behaving primates remain unclear.

*In vitro* studies in brain slices from macaque LPFC, where neurons are disconnected from their networks and their natural inputs are replaced by intracellular currents injected into cell bodies, also report I-SFA (Gonzalez-Burgos et al., 2019; Povysheva et al., 2013; Zaitsev et al., 2012). This raises the fundamental question of whether E-SFA observed during *in vivo* studies could, at least in principle, stem from the intrinsic electrical properties of neurons rather than circuit effects such as the dynamics between excitation and inhibition. The relationship between I-SFA measured *in vitro,* typically in response to constant step-current pulses, and the E-SFA described *in vivo* during behavioral tasks, in which the inputs into a neuron are likely variable and difficult to measure, is not well characterized. Recent work that combined *in vitro* patch clamp and *in vivo* recordings in the electric fish (Akhshi et al., 2023) and mouse (Wei et al., 2023) with computational modelling have explored the contribution of intrinsic mechanisms to heterogeneous spiking activity *in vivo*. Moreover, some studies have integrated I-SFA into computational models of leaky integrate-and-fire neurons (Buonocore et al., 2016; Jolivet et al., 2005; Liu & Wang, 2001) to demonstrate that I-SFA is an important component in replicating activity measured *in vivo*.

In areas that maintain persistent neural activity even in the absence of sensory inputs, such as the LPFC (Fuster & Alexander, 1971; Kubota & Niki, 1971), the interplay between cell-intrinsic dynamics and circuit dynamics is of paramount importance (Mendoza-Halliday & Martinez-Trujillo, 2017; Mendoza-Halliday et al., 2014; Roussy, Mendoza-Halliday, et al., 2021; Torres-Gomez et al., 2020), but remains largely unexplored, especially in primate brains, leaving a critical gap in our understanding of the issue. If the computations that underlie persistent activity landscapes were outsourced to *single neuron intrinsic transfer functions,* synaptic resources could be released to perform other complex operations such as information re-routing that requires complex interplay of excitation-inhibition within local circuits (Yang et al., 2016; Yang et al., 2021). Such area-specific roles of single neuron properties are conceivable, because properties and proportions of cell types vary considerably among neocortical areas (Collins et al., 2010; Condé et al., 1994; Gabbott & Bacon, 1996; Gilman et al., 2017; Kim et al., 2017; Torres-Gomez et al., 2020).

In this study, we combined *in vivo* and *in vitro* electrophysiology in the LPFC of macaque monkeys to study the potential contribution of I-SFA to the neurons’ activity profiles during behavior. First, we recorded the *in vivo* responses of macaque LPFC broad spiking (BS) and narrow spiking (NS), putative excitatory and inhibitory neurons, respectively, during a visually guided saccade task and measured E-SFA. Second, we performed patch clamp experiments in acute macaque LPFC slices and measured I-SFA of responses to step-current inputs in BS and NS neurons. We found that SFA shapes neuronal responses *in vivo* and *in vitro* in both NS and BS neurons. However, in the case of NS neurons, we did not find differences in the decay time constants of I-SFA and E-SFA. By employing a hybrid-modeling approach that involves developing circuit motifs from data-derived Hodgkin-Huxley type models, we finally demonstrated that the lack of such differences in timescales can be attributed to the interplay of inhibitory network dynamics with I-SFA detected *in vitro*.

## Methods

### Experimental Model and Study Participant Details

Two male rhesus macaques (*Macaca mulatta*) were used in both the *in vivo* and *in vitro* recording experiments, ages 10 and 9, weighing 12 and 10 kgs respectively. Additionally, six *Macaca fascicularis* (3 male and 3 female), between 4 and 11 years old and weighing between 5 and 7.3 kgs were used in the *in vitro* experiments. All procedures followed Canadian Council on Animal Care guidelines and were carried out at Western University with approval from the University Animal Care Committee.

### In vivo Recordings

For our *in vivo* experiments, we used two 96-channel Utah arrays (Blackrock Microsystems, Utah, USA) in each monkey, positioned on the dorsal and ventral gyri of the principal sulcus, at the posterior end of the sulcus, targeting area 9/46 (**Fig. 1** top). We targeted layers 2/3 using a shank length of 1.5 mm, and electrode impedances were between 20 and 1500 kΩ. We acquired neural signals at 30 kHz using two 128-channel Cerebus recording systems (Blackrock Microsystems) and saved them for offline sorting. Spike sorting was carried out using Plexon Offline Sorter (Plexon, Texas, USA). Spikes were detected online using a threshold of −3.4 SD of the signal and saved for offline sorting, which was carried out with Plexon Offline Sorter (version 4.5.0, Plexon Inc.). Spike width was calculated as the duration between the trough and peak of the average spike. Units were sorted into either broad spiking (BS), or narrow spiking (NS) based on a threshold calculated as the local minima of the sum of two Gaussians that were fit to the distribution of spike widths (**Fig. S1A**; Torres-Gomez et al. (2020)). A visually guided saccade (VGS) task was used where animals made saccades to a target that appeared on a homogenous background and maintained fixation for 500 ms, at which point the target disappeared and the monkey received reward. We used the onset of the stimulus to identify the neuronal response onset.

**Figure 1.**
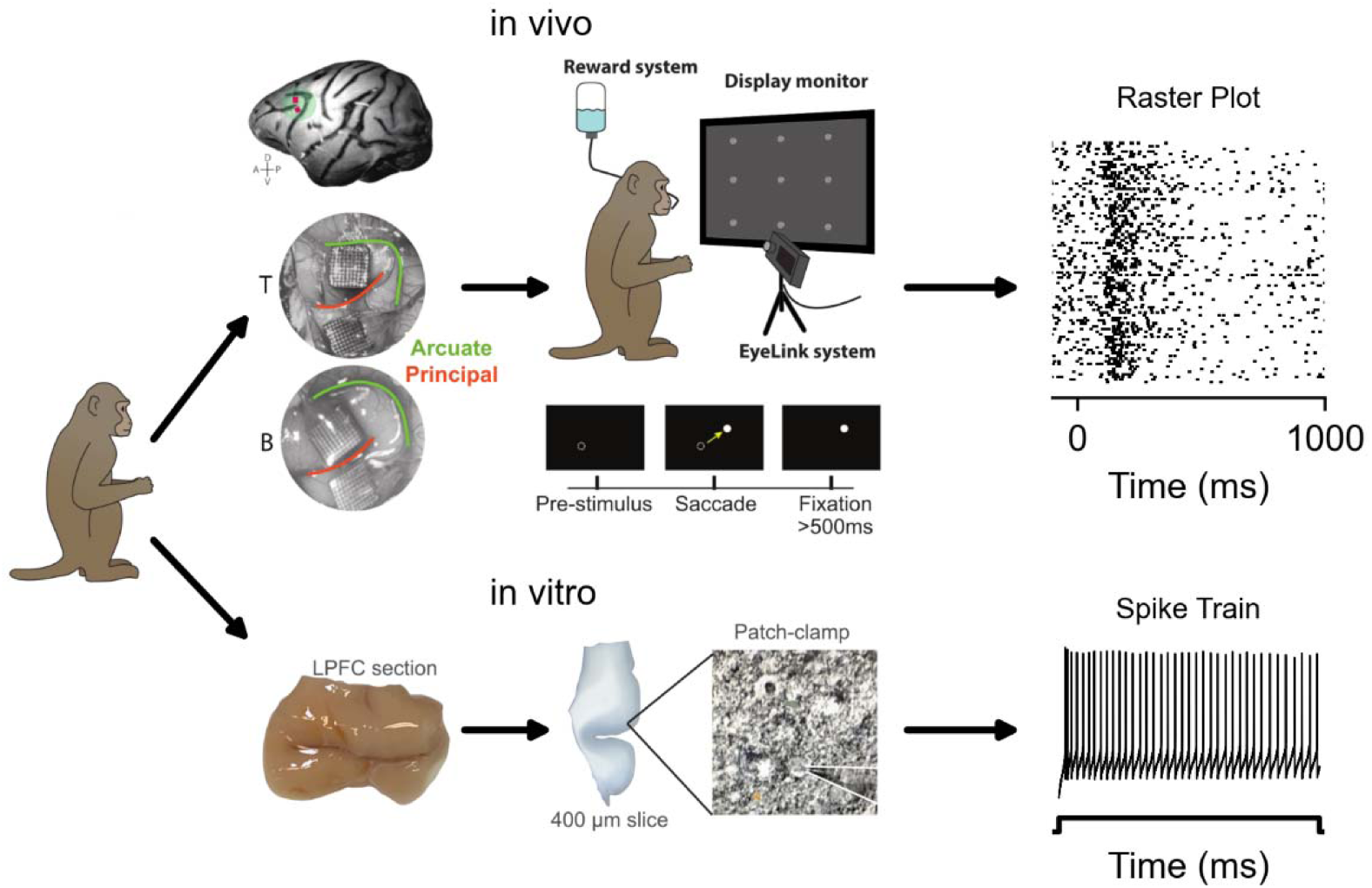
Study design outlining the methods used for neural recordings. Utah arrays were implanted on either side of the principal sulcus, anterior to the arcuate sulcus, in two male rhesus macaques (T and B) and used to record neuronal activity during a VGS task (top). A fixation point appeared on the screen at one of 9 different positions. When the animal fixated, a target appeared at a second position. The animal made a saccade to the target when the fixation point was turned off. Slices from dorsal and ventral banks of the principal sulcus in the LPFC were used for *in vitro* patch clamp recordings (bottom)

### In vitro Preparations

These methods are fully described in Jimenez-Sosa (2020) and briefly summarized here. For the *in vitro* recordings, animals were euthanized, and tissue was collected. Under general anesthesia using isoflurane administration, a craniotomy was performed over LPFC, the dura mater was opened, and then biopsies of LPFC (targeting areas 9/46 and 8a) were taken. Resected tissue was placed in 500 mL of slicing solution (see Jimenez-Sosa (2020)) at 4° C and was maintained at this temperature for approximately 5 minutes of transportation to the neurophysiology lab.

Slices (300 µm thick) were cut perpendicularly to the pial surface (with each slice containing both pia and white matter) using a Leica 1200S vibratome while samples were immersed in ice-cold slicing solution. Slices were placed in slicing solution at 30° C to settle for 15 minutes, and then transferred to chambers with room temperature HEPES solution (see Jimenez-Sosa (2020)) for at least 1 hour before recording. Oxygen levels were maintained at 95% O2 and 5% CO2 throughout this process.

### In vitro Electrophysiological Recordings

Slices were placed in the recording chamber and perfused at 2 mL/min with artificial cerebrospinal fluid (aCSF, see Jimenez-Sosa (2020)) at 34±1° C. The aCSF temperature was continuously monitored and recorded throughout the brain slice experiments. Neurons were visualized for recording under 40X magnification using Dodt gradient contrast system (Dodt et al., 1998) and an infrared camera (Dage-MTI IR-1000).

Recordings were obtained with a MultiClamp 700B amplifier (Molecular devices, California, USA) operating in current clamp mode with an online filter at 10 kHz. The data were digitized at a 20 kHz sampling rate with an Axon Digidata 1440A (Molecular devices, California, USA) and stored via pClamp). Fast glutamatergic and GABAergic synaptic transmission were blocked with 2 mM kynurenic acid and 100 µM picrotoxin to ensure isolation of neurons from local circuitry. Neurons were targeted for recording in all cortical layers. Interneurons were targeted based on having a small cell body with a round or oval shape, and not having an apical dendrite. Pyramidal cells, the most prevalent neurons, were also targeted based on morphology and were well represented in our data. Whole-cell recordings commenced once a seal of 1 GΩ was achieved. Stimuli consisted of one-second current pulses starting at −110 pA and increasing in 20 pA steps to rheobase + 270 pA. The recording sessions could last up to 17 hours, depending on the condition of the slices.

Recordings were analyzed by custom MATLAB scripts that extracted electrical features from the 1 s current step. Activity detection was carried out during the stimulation period, using a minimum threshold of −20 mV. Any sweeps with no activity above this threshold were discarded, along with any sweep with any activity above this threshold before stimulus onset. Sweeps that passed this threshold were then analyzed for spikes using the triangle method of Zack et al. (1977) (**Fig. S1B**) to calculate the threshold for spike detection. The measured voltages during the stimulus were then binned in .5 mV bins from −60 mV to 30 mV to generate a histogram, and then a line was drawn between the peak of the histogram and the bin with the highest values on the right, forming the base of the triangles. A triangle was then formed with each bin value between the two ends of the base, and the perpendicular distance between the base and the bin count was calculated for each bin. The right side of the bin that had the longest perpendicular distance from the base was then used as the critical value. The threshold was then calculated using the following equation

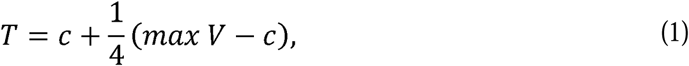

where *T* is the sweep threshold, *c* is the critical value, and *max V* is the maximum voltage across the sweep. This was calculated for every sweep that had a voltage above −20 mV, and the highest threshold for that neuron was used for all sweeps.

To calculate the width of the spikes, we used the distance between the steepest part of the rise and fall by calculating the distance between the peak and trough of the derivative of the voltage. We calculated this for the first 1-3 spikes for each sweep and used the mean.

### Intrinsic and Extrinsic Spike Frequency Adaptation (SFA) Analysis

Adaptation was quantified using an exponential decay function

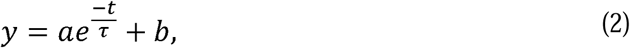

where *a* is the coefficient that scales the adaptation, *b* represents the baseline level to which the adaptation firing rate decays to, and τ is the time constant of adaptation all estimated using custom-made MATLAB scripts. Fitting was limited to minimal τ of 63.2% of the PSTH bin width (6.32 ms for 10 ms *in vivo* bin widths) for E-SFA analysis and to minimally 63.2% of the first ISI for *in vitro* I-SFA analysis. *In vitro* analyses began by separating the cells into BS (n=52) and NS (n=33) based on spike width. Morphology was characterized for about half the recorded cells; hence we used the narrowest spike width among the morphologically confirmed pyramidal cells as our threshold between NS and BS cells to be able to compare between NS and BS cells *in vivo*. A single step-current stimulus close to rheobase capable of producing a train of spikes was chosen for analysis in each neuron based on two criteria: i) at least 4 spikes were fired in response to the stimulus, and ii) the stimulation had to be lower than the greater of either 120 pA, or twice the pA of the rheobase. The instantaneous firing frequency response (reciprocal of the inter-spike intervals (ISIs)) was then calculated and fit from the first spike time onwards using Eq. (2). Additionally, an adaptation index was calculated as the ratio of the first ISI to the last ISI (equivalent to the ratio of the last instantaneous firing frequency to the first), indicating that this index decreases with a greater I-SFA. The latency to first spike was calculated as the time difference between step stimulus onset and the peak of first spike.

For the VGS *in vivo* task, two different types of peristimulus time histograms (PSTHs) were generated: 1. Individual neuron PSTHs (with bin size of 10 ms), where the PSTH is calculated across all trials for a given cell. 2. Population level PSTHs (with bin size of 50 ms), where the PSTH is calculated across the same trial for all neurons in a cell type (n=70 trials for BS and NS). Bin sizes were selected to account for different sample sizes contributing to them while avoiding empty bins. All PSTHs were normalized by the number of trials (individual neuron PSTHs) or cells (population level PSTHs) from which they were calculated, as well as the bin size to obtain PSTHs for the average firing responses per trial or cell, respectively. Individual neurons with post-stimulus response greater than 5 standard deviations away from the pre-stimulus baseline firing rate were considered as selective to the VGS task (BS n=24, NS n =7) and used for further analysis. The exponential decay function of Eq. (2) was fit to the PSTH from the maximum PSTH value onwards. Decay time constants were not different with exponential decay fitting one bin before or after the bin containing the maximal value. Response latency was computed for the VGS task as the time after stimulus presentation for the firing of VGS selective neurons to exhibit a post-stimulus <u>s</u>pike <u>d</u>ensity function (SDF) greater than 5 standard deviations away from the pre-stimulus baseline firing rate; here, SDF was created by convolving the spiking response with an <u>e</u>xcitatory <u>p</u>ost<u>s</u>ynaptic <u>c</u>urrent (EPSC) kernel (with τ*_rise_* = 0.05 ms, τ*_decay_* = 5.3 ms; (Dura-Bernal et al., 2023)). The time to maximal response (also called time to peak response) for the VGS task was computed as the interval between stimulus presentation and peak PSTH response. Adaptation index was calculated as the ratio of the baseline firing rate (b in Eq. (2)) over the peak firing response (*a*+ *b* of Eq. (2)) for the exponential decay function fit to the *in vivo* task 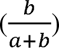.

### Hybrid Circuit Modelling

A Hodgkin-Huxley-type fast-spiking cortical interneuron model by Pospischil et al. (2008) was used to describe NS cells taking into account the more general description of SFA current from Prescott et al. (2006). It is comprised of leak (*I_L_*), Na^+^ (*I_Na_*), delayed rectifier K^+^ (*I_kd_*) and adaptation K^+^ (*I_adapt_*) currents. A population of n=35 NS models was generated by fitting each model’s I-SFA time constant (dictated by τ*_max_* of the *I_adapt_* current activation, **Table S1**) during a 3µA/cm^2^ step input current to the time constant of I-SFA obtained *in vitro* from slice recordings of NS neurons. All other model parameter values were taken from (Pospischil et al., 2008): L = d = 56.9 µm, *g_leak_* = 3.8 × 10^−5^ S/cm^2^, *E_leak_*= −70.4 mV, 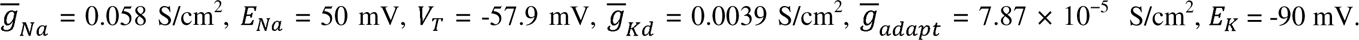

*In vivo* spiking data from BS neurons across 70 trials were pooled to create a dataset consisting of 70 trials with each trial containing spiking activity from a population of BS neurons (n=24). Data was pooled across animals as no discernable differences were observed in exponential decay properties or maximal response latencies among them (**Fig S2**). The combined spike train of each trial was then convolved with EPSC kernel as before (Dura-Bernal et al. (2023)) to generate a SDF that was used as the excitatory input into the population of models. These SDFs were causally low-pass filtered (50 Hz cutoff) and convolved with an inhibitory <u>p</u>ost<u>s</u>ynaptic <u>c</u>urrent (IPSC) kernel (with τ_rise_= 0.07 ms, τ_decay_= 18.2 ms; Dura-Bernal et al. (2023)) to generate the feedforward inhibitory input. Each model then received the excitatory and inhibitory inputs (70 trials total) to examine the impact of feedforward inhibition on NS neurons. The PSTH of the resulting spiking outputs of NS neurons, receiving either solely excitatory or mixed inputs, were generated for each model and an exponential decay was fit as described above for the *in vivo* data (see *Intrinsic and Extrinsic Spike Frequency Adaptation (SFA) Analysis*).

Synaptic bombardment was implemented as Ornstein-Uhlenbeck processes following (Destexhe et al., 2001) with g_e0_ = 0.0320 mS/cm^2^, Θ_e_ = 0.2516 ms^-1^, μ_e_ = 0.0320 mS/cm^2^, σ_e_ = 0.0051 mS/cm^2^, g_i0_ = 0.0365 mS/cm^2^, Θ_i_ = 0.1117 ms^-1^, μ_i_ = 0.1827 mS/cm^2^, σ_i_ = 0.0095 mS/cm^2^ and simulated using a stability-optimized adaptive strong order 1.5 and weak order 2.0 method (Rackauckas & Nie, 2018). The modelling approach did not account for the possibility that feedforward inhibitory cells generating the feedforward inhibition (IoI) possess I-SFA themselves. To address this possibility, we added the high-pass properties of I-SFA (Benda, 2021; Benda & Hennig, 2008; Benda & Herz, 2003) to the low-pass filter feedforward inhibitory component at 3 different high-pass cutoffs (0.1 Hz, 3 Hz and 5 Hz) within the range of I-SFA high-pass cutoffs (Benda & Hennig, 2008). Simulation and analysis methods for the mixed input with band-pass properties were otherwise unchanged from the low-pass only mixed input simulations.

### Numerical Methods and Software

All simulations were performed using the Julia programming language (Julia v1.10.1; Bezanson et al. (2017)), utilizing an adaptive order quasi-constant timestep numerical differentiation function method (QNDF) implemented in DifferentialEquations.jl (Rackauckas & Nie, 2017). Fitting of model I-SFA time constant was done using an adaptive differential evolution (DE/rand/1/bin) with radius limited sampling in Optimization.jl (Dixit & Rackauckas, 2023). Figures were generated using matplotlib 3.4.3 and seaborn 0.11.2 in Python 3.10. The code for reproducing the modelling results and figures is available online using the link: https://github.com/nkoch1/LPFC_adaptation.git

### Quantification and Statistical Analysis

Statistical significance was set at *p*<0.05, with the exact *p* and n values reported in the Results Section. Two-way comparisons of differences were computed using the Mann-Whitney-Wilcoxon two-sided test (scipy 1.11.1 in Python). Three(and more)-way comparisons were done with a Kruskal-Wallis test, with significant outcomes followed by post-hoc analysis performed with Dunn’s test (scipy 1.11.1 in Python). Data are reported as median (interquartile range).

## Results

### In vivo Extrinsic Adaptation (E-SFA) in BS and NS units

We recorded the responses of 325 LPFC neurons in areas 9/46 of two macaque monkeys performing a VGS task using MEAs (Utah arrays, **Fig. 1** top). During the task, a fixation point appeared at one of nine positions on the screen; when the animal fixated on the stimulus, a second stimulus appeared at a different position and the animals made a saccade toward the new stimulus (**Fig. 1** top). We sorted single units and obtained average waveforms as well as raster plots of spike events. We divided the units into broad (BS) and narrow spiking (NS) according to the trough-to-peak distribution of their waveforms (**Fig. 2A-B** see Methods). After the target onset, many of the area 9/46 units (BS n =24; NS n =7) showed a ‘visual response’ in agreement with findings from previous studies of LPFC area 8A (Bullock et al., 2017). The PSTHs of BS and NS units were computed (**Fig. 2C-E**) and fit with an exponential decay function of Eq. (2) from their post-stimulus peak. The PSTHs of BS and NS units were aligned to the target stimulus presentation (**Fig. 2D-E** middle), as well as aligned to the time of maximal response and normalized by this maximal to better demonstrate the decay kinetics of the responses (**Fig. 2D-E** bottom). This post-stimulus time to maximal response (T_Max_) was longer in the BS units (median 300 ms (IQR 190-395 ms)) compared to NS units (median 145 ms (IQR 130.0-237.5 ms); **Fig. 2F**; Mann-Whitney-Wilcoxon test two-sided, U=175.5, p=0.016) with maximal BS unit responses occurring up to 600 ms after stimulus presentation. This latter result could be attributed to the heterogeneity of inputs into BS pyramidal cells within the LPFC as well as varying contrast sensitivities that influence response latencies (Bullock et al., 2017). No differences in the scaling coefficient (a in Eq. (2); **Fig. 2G**) were detected between BS (median 17.869 Hz (IQR 11.331-25.977 Hz)) and NS (median 17.798 Hz (IQR 15.871-56.666 Hz)) units (Mann-Whitney-Wilcoxon test two-sided, U=58.0, p=0.234). The baseline firing rate (b in Eq. (2); **Fig. 2H**) of the exponential decay was also not significantly altered (Mann-Whitney-Wilcoxon test two-sided, U=57.0, p=0.216) between BS (median 2.033 Hz (IQR 0.000-5.859 Hz)) and NS (median 5.625 Hz (IQR 2.435-9.950 Hz)) units.

**Figure 2.**
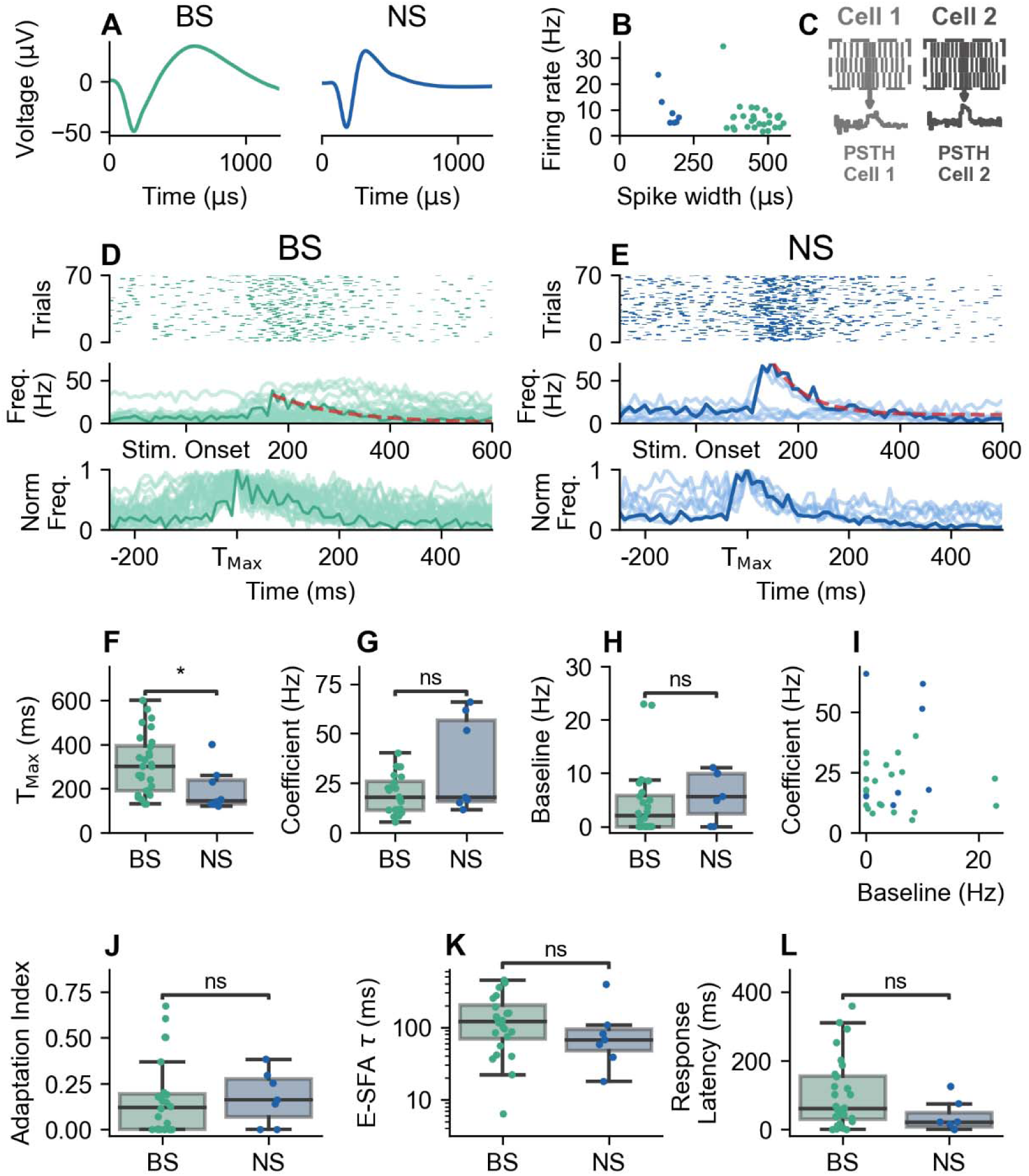
*In vivo* adaptation for VGS task. (A) Average waveform for an example BS unit (left) and an example NS (right) unit. (B) Spike width vs firing rate for responsive units, labelled as either NS or BS units. (C) PSTHs generated per cell for (D) BS and (E) NS neurons - with example exponential decay function fits (red dashed lines) - aligned to stimulus onset (Stim. Onset, middle) as well as the PSTHs normalized by the peak firing rate (Norm F) aligned to maximal response time (T_Max_; bottom). (F) Post-stimulus time to maximal PSTH responses (T_Max_) in both BS and NS units. (G, H) Comparison of the coefficients (G), and baselines (H) of adaptation between BS and NS units, obtained through fitting th exponential decay function of Eq. (2) to the decay phases of their PSTHs (red lines in d and e). (I) Relationship between the coefficients and baselines of adaptation (highlighted in G and H). (J) Adaptation indices, (K) decay time constants obtained through fitting the exponential decay function of Eq. (2), and (l) post-stimulus response latencies in both BS and NS units. ns p≥0.05, * p < 0.05, **p<0.01, *** p < 0.001

The scaling coefficient and the baseline firing did not have monotonic relationships in BS (Spearman ρ = −0.187, p = 0.381) or NS (Spearman ρ = 0.286, p = 0.535) cells (**Fig. 2I**). The magnitude of *in vivo* E-SFA, quantified by the adaptation index, did not differ between BS (median 0.120 (IQR 0.000, 0.193)) and NS (median 0.161 (IQR 0.070, 0.274)) units (Mann-Whitney-Wilcoxon test two-sided, U=72, p =0.595; **Fig. 2J**). Crucially, we did not detect any difference in the time constants of decay between the populations of BS (median 120.206 ms (IQR 69.631-205.942 ms)) and NS (median 67.219 ms (IQR 46.170-94.029 ms)) units (**Fig. 2K**; Mann-Whitney-Wilcoxon test two-sided, U=113.0, p=0.182). Taken together, these results indicate that E-SFA is generally similar between BS and NS units. However, the latency to the maximal response is longer on BS than in NS units. Given that BS units serve as inputs to NS units (Lewis et al., 2002; Melchitzky et al., 2001; Melchitzky & Lewis, 2003), this finding may suggest a more precise regulation of stimulus timing within NS units even though there is no significant difference in response latency between BS (median 60.563 ms (IQR 30.117-155.172 ms) and NS (median 22.042 ms (IQR 8.257-48.697 ms)) units (**Fig. 2**l; Mann-Whitney-Wilcoxon test two-sided, U=14.0, p=0.08).

### Circuit Dynamics during VGS Task Accentuate Disparities between BS and NS Responses in vivo vs in vitro

To obtain an estimate of the population responses *in vivo* during the VGS task, we pooled across BS and across NS cells per trial to obtain population response estimates for BS and NS neurons, normalized by the population size. We examined the average population response (**Fig. 3A**) to a given trial of the VGS task by taking a population-wide PSTH for 70 trials (**Fig. 3B-C** for BS and NS units, respectively). These population-wide PSTHs were aligned to stimulus onset (**Fig 3B-C** top) as well as to the time of maximal population response and normalized by this maximal response (**Fig 3B-C** bottom). This post-stimulus time to maximal population response (T_Max_) was longer in the BS population (median 300 ms (IQR 250-350 ms)) than in the NS population (median 150 ms (IQR 150-200 ms); **Fig. 3D**; Mann-Whitney-Wilcoxon test two-sided, U=4290, p=7.74*10^-19^). Fitting these population-wide responses from their post-stimulus peak with the exponential decay function of Eq. (2) revealed that the NS population responses have larger magnitude (coefficient) of decay (median 25.521 Hz (IQR 20.717-31.770 Hz)) than the BS population (median 12.180 Hz (IQR 10.434-13.570 Hz); **Fig. 3E**; Mann-Whitney-Wilcoxon test two-sided, U=8.0, p=2.70*10^-22^), and that the average firing rate is higher in NS (median 7.472 Hz (IQR 6.231-8.440 Hz)) than BS (median 4.524 Hz (IQR 2.461-5.430 Hz)) populations (**Fig. 3F**; Whitney-Wilcoxon test two-sided, U=403.0, p=9.97*10^-17^). Interestingly, when comparing the baseline to the coefficient of the exponential decay, there is distinct clustering of narrow and broad population responses (**Fig. 3G**) that is not evident in the individual responses *in vivo* (**Fig. 2I**) or *in vitro* (**Fig. 4G**); the negative relationship found *in vitro* (but not *in vivo*) was present in the population level in BS (Spearman ρ = - 0.423, p = 0.0003) but not in NS (Spearman ρ = −0.233, p = 0.057), suggesting that the response dynamics of these neurons are less influenced by circuit effects in a manner that needs to be further investigated.

**Figure 3.**
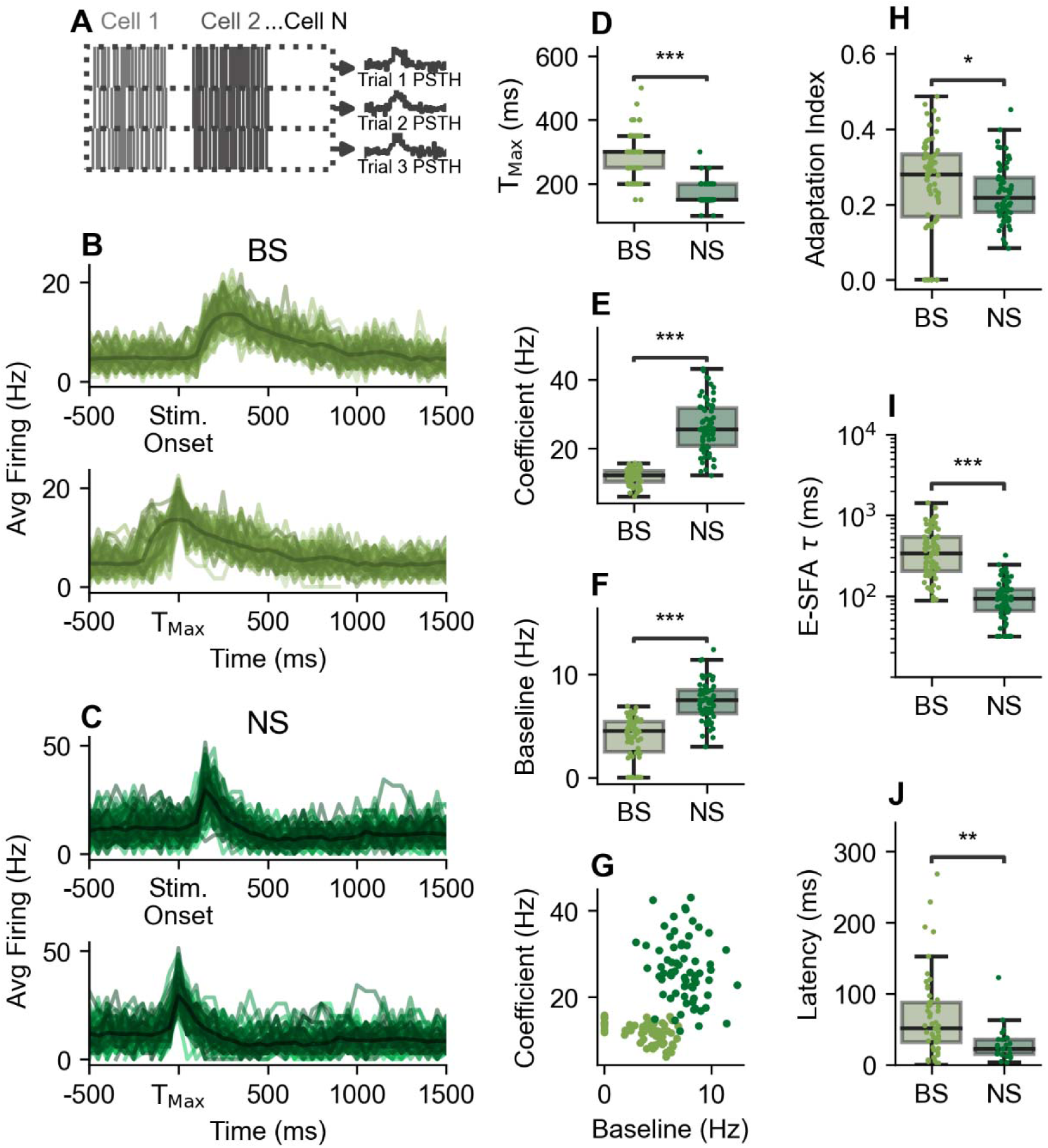
*In vivo* adaptation at the population level. (A) Illustration of how population level PSTHs are computed. Cell responses were pooled across the population of BS or NS units for each VGS task trial. (b, c) The computed population level PSTHs for BS (B) and NS units (C) aligned to stimulus onset (top) and to trial maximal population response time (T_Max;_ bottom). (D) Post-stimulus time to maximal PSTH responses (T_Max_) in both BS and NS population responses. (E, F) Comparison of the coefficients (E), and baselines (F) of adaptation between BS and NS population responses, obtained through fitting th exponential decay function of Eq. (2) to the decay phases of their population level PSTHs (highlighted in B and C). (G) Relationship between the coefficients and baselines of adaptation (highlighted in E and F). (H) Adaptation indices, (I) decay time constants obtained through fitting the exponential decay function of Eq. (2), and (J) Post-stimulus response latencies of the population level BS and NS responses. ns p≥0.05, * p < 0.05, **p<0.01, *** p < 0.001

**Figure 4.**
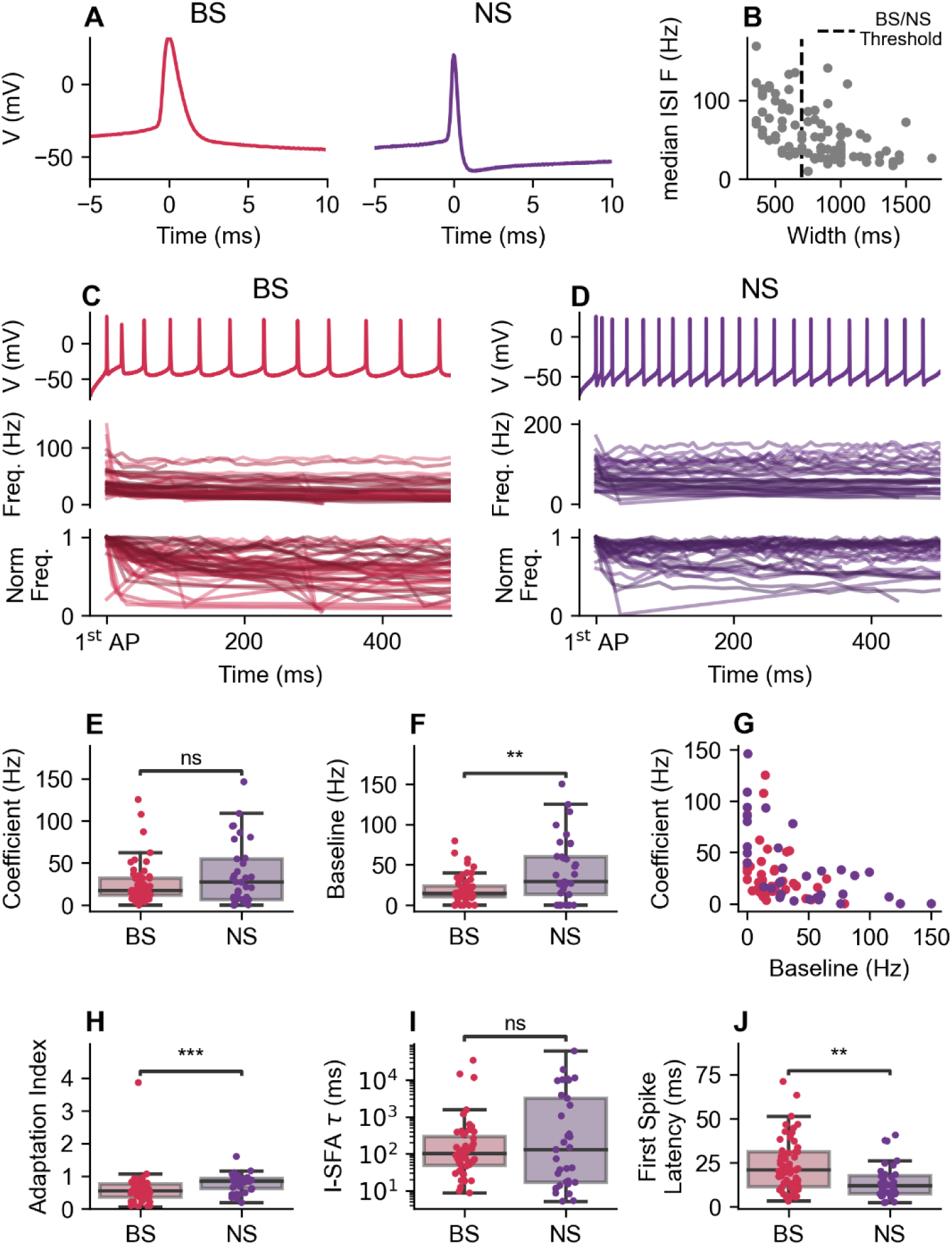
*In vitro* slice recording SFA. (A) Spike shape for an example BS (left) and an example NS (right) neuron. (B) Distribution of neuron spike width and frequency response of median ISI in relation to the BS/NS threshold. (C, D) Sample recordings of a BS (C) and NS (D) neuron (top) along with th profiles of the firing frequency (F) and frequency normalized by the peak firing rate (Norm F) of all recorded (C) BS and (D) NS neurons (bottom). (E, F) Comparison of the (E) coefficients, and (F) baselines of SFA between BS and NS neurons, obtained through fitting the exponential decay function of Eq. (2) to the profiles of the firing frequency (in C and D). (G) Relationship between the coefficients and baselines of SFA (highlighted in E and F). (H) Adaptation indices, (I) decay time constants obtained through fitting the exponential decay function of Eq. (2), and (J) latencies between the start of the step-current stimulus and peak of first spike for both BS and NS neurons. ns p≥0.05, * p < 0.05, **p<0.01, *** p < 0.001

By comparing the adaptation index between the BS (median 0.279 (IQR 0.167-0.333)) and NS (median 0.218 (IQR 0.179-0.271)) population responses, the former was found to be larger (**Fig. 3H**; Mann-Whitney-Wilcoxon test two-sided, U=2778, p=0.043). Notably, the variability in individual BS latencies (**Fig. 2L**) resulted in larger population BS response decay time constants (median 339.492 ms (IQR 201.873-538.272 ms)) compared to those of the NS population (median 91.932 ms (IQR 65.594-120.747 ms); **Fig. 3I**; Mann-Whitney-Wilcoxon test two-sided, U=4339, p=1.11*10^-18^). In contrast, the response latency in BS (median 51.086 ms (IQR 31.826-88.096 ms) population was slower than in NS (median 22.458 ms (IQR 14.588-35.603 ms)) population (**Fig. 3J**, Mann-Whitney-Wilcoxon test two-sided, U=86.3, p=0.0013), further highlighting that circuit dynamics may exacerbate differences between BS and NS responses *in vivo* compared to *in vitro*.

### Intrinsic Spike Frequency Adaptation (I-SFA) Mostly Differs between BS and NS Neurons

To explore the *in vitro* I-SFA in individual neurons isolated from the network, we conducted patch clamp recordings in LPFC slices of 8 animals (see methods and **Fig. 1** bottom). Neurons were classified into BS (n=53) and NS (n=33) depending on a threshold for the width of the action potentials, defined based on the spike waveform shape and the median of their interspike interval (ISI) distribution (**Fig. 4A-B**). Comparing the *in vitro* firing responses of LPFC BS and NS cells to injected step-currents (**Fig. 4C-D**) revealed that the scaling coefficient of the exponential decay is not different between BS (median 17.115 Hz (IQR 11.734-31.974 Hz)) and NS (median 27.084 Hz (IQR 6.660-54.819Hz)) cells (**Fig. 4E**; Mann-Whitney-Wilcoxon test two-sided, U=801.0, p=0.61). It also revealed that the baseline of the exponential decay is larger for NS (median 29.004 Hz (IQR 13.181-60.330 Hz)) than BS (median 14.591 Hz (IQR 10.428-25.863 Hz)) cells (**Fig. 4F**; Mann-Whitney-Wilcoxon test two-sided, U=567.0, p=0.009). Notably, the relationship between the baseline firing rate and the coefficient of the exponential decay (**Fig. 4G**) although qualitatively similar to those of the neurons recorded *in vivo* (**Fig. 2H**), are negative for BS (Spearman ρ= −0.255, p = 0.068) and NS (Spearman ρ = −0.678, p = 1.467*10^-5^). The adaptation index of BS cells (median 0.545 (IQR 0.365-0.754)) was found to be smaller (indicating more I-SFA) than that of NS (median 0.838 (IQR 0.620-0.935); **Fig. 4H**; Mann-Whitney-Wilcoxon test two-sided, U=456.0 p=0.0003). Fitting the exponential decay function of Eq. (2) to the responses of the BS and NS cells showed that whereas BS decay time constants (median 101.003 ms (IQR 48.710-292.547 ms)) are not different from the NS decay time constants (median 129.364 ms (IQR 16.715-3166.415 ms); **Fig. 4I**; Mann-Whitney-Wilcoxon test two-sided, U=849.0, p=0.949), the BS latency to first spike (median 20.7 ms (IQR 11.200-31.000 ms)) after step-current onset is longer than NS cells (median 11.75 ms (IQR 7.450-17.700 ms); **Fig. 4J**; Mann-Whitney-Wilcoxon test two-sided, U=647.5, p=0.0014). The latter suggests that NS neurons undergo fast adaptation and then stabilize their firing at a certain level as the current stimulus is maintained. The notion that NS neurons adapt to a high basal firing rate is also evident in the large adaptation index of these neurons (**Fig. 4H**). Importantly, NS neurons have been often described as linear filters showing almost no I-SFA (Ascoli et al., 2008). These findings thus indicate that at least some NS LPFC neurons show non-linear response profiles due to I-SFA early after stimulation *in vitro*.

### LPFC BS and NS Neurons Exhibit Larger but not Faster Adaptation in vivo than in vitro

The magnitude of adaptation for both I-SFA and E-SFA, quantified using adaptation index, was found to be significantly larger (indicating smaller magnitudes of adaptation) *in vitro* than *in vivo (***Fig. 5A-D**; Kruskal Wallis H-test, H=53.51, p=1.43*10^-11^). Specifically, the adaptation index in both BS and NS neurons *in vitro* was larger than in BS (Dunn’s test, p=6.46*10^-10^ and p =6.11*10^-10^ respectively) and NS (Dunn’s test, p=0.006 and p=0.0002 respectively) *in vivo* during the VGS task, suggesting that the magnitude of adaptation is greater *in vivo* than *in vitro.* However, the decay time constants of the BS and NS firing response *in vivo* were not different from those obtained with *in vitro* patch clamp step-current protocols except for the slower BS population responses (**Fig. 5E-H**; Kruskal Wallis H-test, H=60.56, p=9.31*10^-12^). It is important to note, however, that the variance in the NS firing response decay time constants *in vitro* was far more pronounced than those *in vitro*.

**Figure 5.**
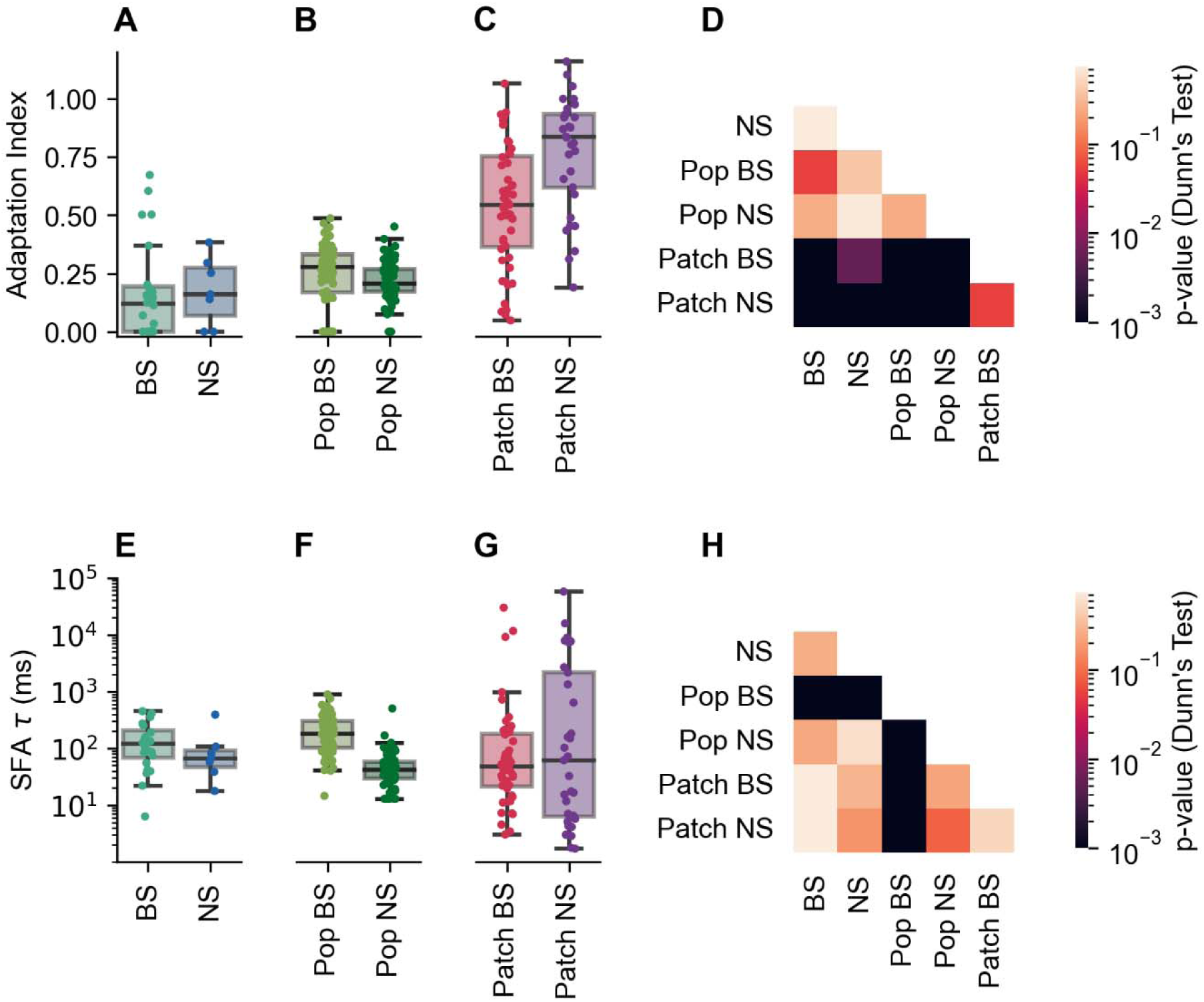
Individual neuron SFA is not different between *in vivo* VGS task and *in vitro* step protocol (Patch) nor across cell type (BS/NS). (A) Adaptation index of each BS and NS unit during the VGS task. (B) Adaptation index of the population of BS and NS units during each VGS trial (Pop BS/NS). (C) Adaptation index of BS and NS neurons in vitro in response to a step-current (Patch BS/NS). (D) The p-values of Dunn’s post hoc test for the adaptation index after a Kruskal-Wallis test (H=53.51, p=1.43*10^-11^). (E-G) Fitted exponential decay time constants obtained from Eq. (2) for (E) the PSTH of each BS and NS unit across trials (BS/NS), (F) the PSTH of the population of BS and NS units (Pop BS/NS), and (G) the frequency responses of BS and NS neurons to a step-current (Patch BS/NS). (H) The p-values of Dunn’s post hoc test for the time constants after a Kruskal-Wallis test (H=60.56, p=9.31*10^-12^). ns p≥0.05, * p < 0.05, **p<0.01, *** p < 0.001

### Excitation and Feedforward Inhibition of Inhibition in a Hybrid Circuit Model Bridges the Gap between in vitro I-SFA and in vivo E-SFA

To understand the link between I-SFA, estimated *in vitro,* and *in vivo* E-SFA of responses during the VGS task, we implemented a hybrid approach in which a spike density function (SDF) generated using the *in vivo* recorded BS firing activity was used as the input into a population of NS Hodgkin-Huxley model neurons. The circuit model reasonably assumed that NS interneurons are recruited mostly by excitatory input provided locally by BS pyramidal cells. In this hybrid approach, it was assumed implicitly that (i) the *in vivo* BS firing activity is already accounting for all the inputs and feedback these cells receive from other cell types including NS neurons, and (ii) the NS population consists of model neurons with the same I-SFA time constant as those fit to the *in vitro* recordings (**Fig. 4D** and **I**). The response of the population of NS models to this ‘realistic’ *in vivo* excitatory input was broad and had a slow decay in their firing rate (**Fig. 6A**). Predictably, this response is not qualitatively similar to either the individual (**Fig. 2E**) or population (**Fig. 3C**) responses of NS cells recorded *in vivo*.

**Figure 6.**
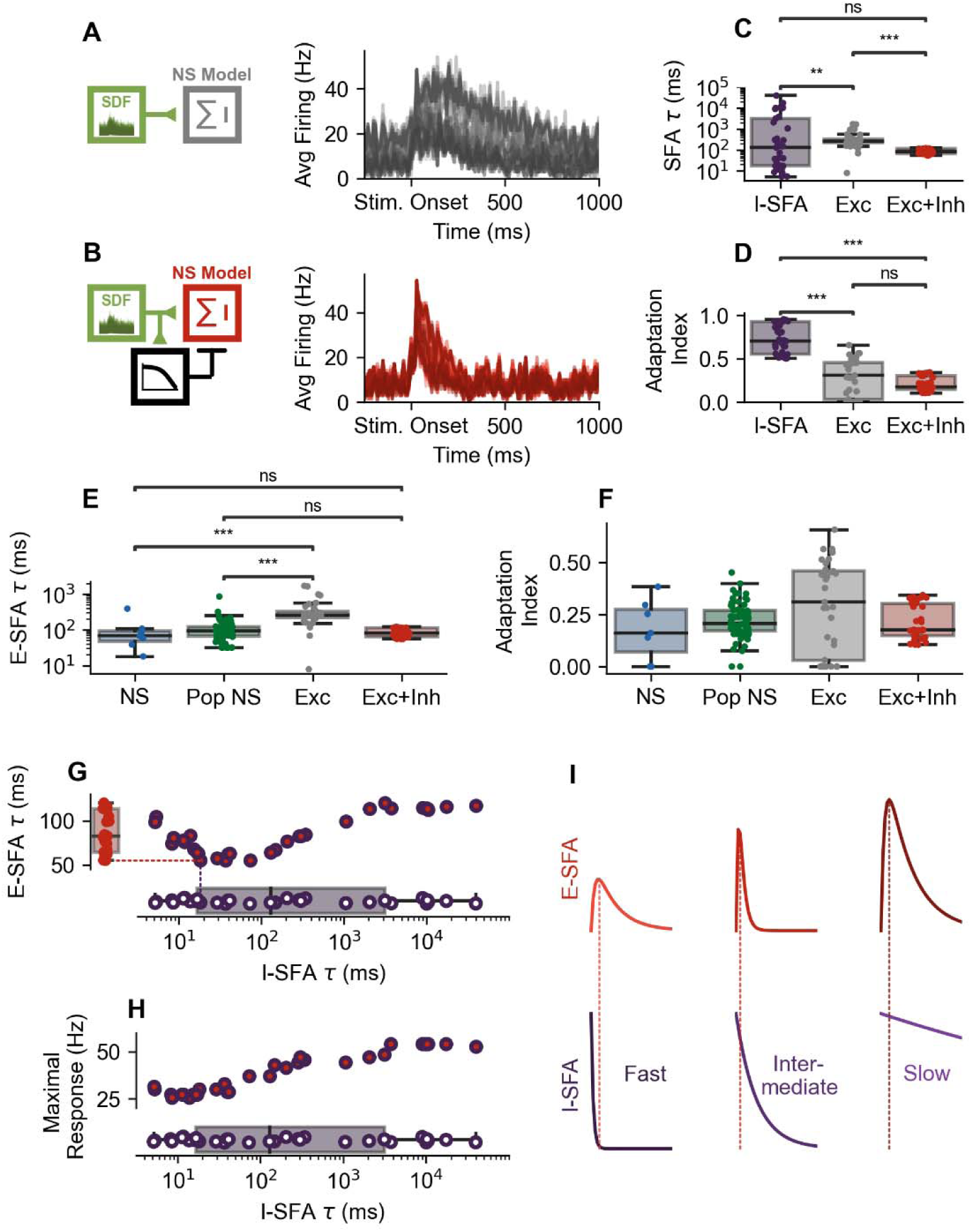
Responses of a population of NS model neurons to BS input and feedforward inhibition. (A) Left: schematic depiction of the hybrid circuit model wherein *in vivo* data from BS units (spike density function (SDF); green) was injected as the excitatory input into a population of NS model neurons (grey). Right: the computed PSTHs generated by these NS model neurons. (B) Left: schematic depiction of the hybrid circuit model wherein *in vivo* data from BS units (green) was injected as the excitatory input to a population of NS model neurons (red) which concurrently receives feedforward inhibition (black). Right: the computed PSTHs generated by these NS model neurons. (C) The exponential decay time constants (computed by fitting Eq. (2)) of the responses of the population of NS model neurons to the step input, BS input and BS input with feedforward inhibition (Kruskal Wallis H-test, H=21.92, p= 1.69*10^-5^). (D) The adaptation index for the step input, BS input and BS input with feedforward inhibition (Kruskal Wallis H-test, H=61.02, p=5.62*10^-14^. (E) Comparison between experimental (NS, Pop NS) and model (BS, BS+Inh) time constants computed by fitting Eq. (2) (Kruskal Wallis H-test, H=52.15, p=2.83*10^-14^), and (F) adaptation index (Kruskal Wallis H-test, H=5.91, p=0.12). Dunn’s post hoc test p-values are reported after a Kruskal-Wallis tests. (G) The relationship between I-SFA (purple; 3µA/cm^2^ step) and the E-SFA decay time constant (red) of feedforward inhibition (obtained in b). (H) The relationship between I-SFA (step input) and the maximal response. (I) Schematic illustrating the decrease in E-SFA time constant in NS models with intermediate I-SFA time constants in response to the circuit input. ns p≥0.05, * p < 0.05, **p<0.01, *** p < 0.001.

To approximate the *in vivo* response of the NS cells, we added feedforward inhibition to the inputs received by the NS model neurons. This feedforward inhibitory component consisted of low-pass filtered excitatory input from BS responses that was subsequently convolved with an IPSC kernel. The population of NS model neurons receiving this mix of excitatory inputs from BS neurons and the new feedforward inhibitory inputs (Inhibition of Inhibition or IoI) exhibited a response with a faster decay (**Fig. 6B**), resembling the individual and population NS responses recorded *in vivo*.

We compared the I-SFA decay time constants of this population of NS model neurons receiving a step-current (median 129.370 ms (IQR 16.704-3142.149 ms)) to the E-SFA decay time constants of the same population of model neurons receiving the data-driven excitatory BS input (median 254.761 ms (IQR 217.756-328.491 ms)) as well as the mixed excitatory/inhibitory input (median 82.308 ms (IQR 64.240-113.468 ms); **Fig. 6C**). Our results revealed that differences exist in the responses of the population of model neurons to step, excitatory and mixed inputs (Kruskal Wallis H-test, H=21.92 p=1.69*10^-5^), with the responses to excitatory input having decay timescales different from the step input (Dunn’s test, p=0.0067) and from the mixed input (Dunn’s test, p=3.1 *10^-6^). Additionally, the I-SFA of the population of NS models (i.e. the response to step-current input) was not different from the E-SFA decay timescales of the population of models to the mixed input (Dunn’s test t, p=0.051). This is in line with the lack of differences in the decay timescale of the experimentally obtained NS I-SFA and VGS response decay and the differences in the variance (**Fig. 5E-H**). Importantly, the *in vivo* timescales of E-SFA were not achieved by synaptic bombardment (Destexhe et al., 2001; Destexhe et al., 2003; Paré et al., 1998) in the absence of feedforward IoI (**Fig. S3**).

The magnitude of adaptation, quantified using the adaptation index, of model responses to step-current input (median 0.702 (IQR 0.553-0.928)), data-driven excitatory input (median 0.310 (IQR 0.031-0.458)) and mixed excitatory/inhibitory input (median 0.177 (IQR 0.148-0.302)) were also found to be different between the three different conditions (Kruskal Wallis H-test, H=61.02, p=5.62*10^-14^; **Fig. 6D**). The step-current adaptation indices were larger (indicating smaller magnitudes of adaptation) than the ones resulting from the data-driven excitatory input (Dunn’s test, p=9.06*10^-10^) and the adaptation indices resulting from the mixed excitatory/inhibitory input (Dunn’s test, p= 3.84*10^-11^). Furthermore, the magnitude of adaptation was not different with the addition of inhibition to the circuit (Dunn’s test, p=0.26) compared to excitation alone.

The time constants of the responses of the model neurons were additionally compared to the time constants of NS neurons receiving the excitatory and mixed inputs, and their individual and population responses *in vivo* (**Fig. 6E**; Kruskal Wallis H-test, H=52.12, p=2.83*10^-11^). The model time constants in response to the excitatory input differed from the *in vivo* NS (Dunn’s test, p=4.81*10^-5^) and population NS (Dunn’s test, p=1.41*10^-10^) time constants, whereas the time constants of the population of models with the mixed input was not statistically different from the *in vivo* NS (Dunn’s test, p=0.50) and population NS (Dunn’s test, p=0.64). Additionally, by comparing adaptation indices of model neurons in response to the excitatory and mixed inputs to the *in vivo* individual NS and population-level NS values (**Fig. 6F**; Kruskal Wallis H-test, H=4.24, p=0.24), we found that the magnitude of E-SFA of model neurons to the excitatory and mixed inputs are similar to the *in vivo* NS and the population NS responses. It is important to note that the modelling approach used here was agnostic to the cellular identity and composition of the neurons that provide IoI. Despite this agnosticism, the modelling approach did not account for the possibility that feedforward inhibitory cells generating the feedforward inhibition (IoI) possess I-SFA themselves. To explore this possibility, we incorporated the high-pass filtering properties of I-SFA (Benda, 2021; Benda & Hennig, 2008; Benda & Herz, 2003) into the feedforward inhibitory component at 3 different high-pass cutoffs; by doing do, we found that this modification did not alter outcomes or our conclusions **(Fig. S4),** suggesting that whether the inhibitory neurons involved in the feedforward inhibition possess I-SFA has little consequence.

Since the I-SFA time constants of the models are known - unlike the case for *in vivo* recordings – and that the same set of NS models were used in the *in vitro* and the mixed input *in vivo* conditions, we further investigated how the response of the NS models depend on their I-SFA properties. This was done by plotting the time constants of E-SFA of NS model neurons, receiving the data-driven data mixed inputs, with respect to the I-SFA ones; doing so revealed that, at lower levels of I-SFA (as measured by the time constant of the step-current evoked model responses), the decay timescale of the responses of the NS models to the mixed input exhibits biphasic dependence at lower levels of I-SFA (**Fig. 6G**). Specifically, the E-SFA time constants decreased as the I-SFA timescale increased from small I-SFA (i.e., fast NS models; left of **Fig. 6G**) to intermediate I-SFA (middle of **Fig. 6G**) time constants, and then increased again at larger I-SFA time constants (i.e., slow NS models; right of **Fig. 6G**). Curiously, as the time constant of I-SFA further increased, no dependency of E-SFA on I-SFA time constants was observed, indicating that I-SFA is not contributing to the decay in the *in vivo* response (E-SFA). This is reflected in the increasing relationship between the maximal response *in vivo* and the I-SFA time constants (Spearman ρ = 0.915, p = 9.70 *10^-14^; **Fig. 6H**). When I-SFA is fast, the neuron adapts prior to the peak of the embedded response resulting in a lower maximal response (**Fig. 6H** and **I** left). At intermediate timescales of I-SFA, when the I-SFA of the NS neuron is of similar timescale to the decay of the overall circuit input it receives, the interaction of these 2 processes results in a faster decay and a smaller E-SFA time constant (**Fig 6I** middle). In neurons with slow I-SFA, the contribution of I-SFA to E-SFA decay is limited by the relative lack of adaptation during E-SFA resulting in larger maximal response amplitudes (**Fig 6H** and **I** right). Taken together, these results suggest that the contribution of I-SFA to E-SFA is timescale dependent despite the impact of I-SFA on maximal response amplitudes across I-SFA timescales.

## Discussion

In this study, we explored SFA of BS and NS neurons in the LPFC *in vivo* (during a VGS task) and *in vitro*. We found that adaptation is present in both conditions *in vivo* (E-SFA) and *in vitro* (I-SFA) conditions. Firing rates decayed as a function of stimulus onset in both BS and NS neurons. The timescale of adaptation was similar across all cell types and conditions; however, NS cells showed a higher firing rate during *in vitro* recordings compared to *in vivo*. To understand the relationship between E-SFA measured *in vivo* and I-SFA measured *in vitro*, a hybrid approach was used, in which NS cells were described in terms of Hodgkin-Huxley models with I-SFA time constants obtained from fitting *in vitro* data, and then subjected to input from recorded data. Simulations with this hybrid approach suggest that both local feedforward inhibition and I-SFA contribute to the response dynamics observed during the VGS task with contribution of I-SFA to E-SFA in each NS neuron dependent on its I-SFA properties.

### Differences in Activity and Adaptation in vitro and in vivo

Our results showed that the magnitude of *in vitro* I-SFA is larger in BS than NS neurons with step-current stimulation, as quantified by lower adaptation index values (**Fig. 4H**), in line with previous studies of macaque LPFC and rodent cortex slices reporting that BS pyramidal neurons exhibit greater adaptation than NS parvalbumin-positive cells (González-Burgos et al., 2005; McCormick et al., 1985; Povysheva et al., 2013). However, the difference in adaptation magnitude between BS and NS is diminished *in vivo* during the VGS task, and the adaptation index values recorded during the VGS task are notably smaller in both NS and BS neurons than in response to step input stimuli *in vitro* (**Fig. 5A-C**). This is in line with previously reported increases in adaptation magnitude *in vivo* compared to *in vitro (Fernandez et al., 2018)*. This discrepancy between *in vitro* and *in vivo* adaptation magnitudes could stem from differences in the neuronal input across these two recording paradigms. Step stimuli can effectively be used to uncover intrinsic neuronal dynamics (Druckmann et al., 2011), but may be limited in their generalizability. For example, the impact of plasticity in neuronal ion channel composition on firing rate is dependent on whether static inputs (such as current steps) or dynamic inputs (i.e. synaptic inputs) are used to probe intrinsic neuronal excitability (Szücs & Huerta, 2014). Furthermore, rheobase and spike counts obtained from step-current stimuli have poor predictive power of neuronal responses to simulated synaptic inputs especially at low firing rates (Szabó et al., 2021). These limitations associated with using step-current stimuli may extend to not only I-SFA but also to E-SFA. To address this, we employed a hybrid approach in which data-driven synaptic inputs were embedded in models parameterized by step-current evoked I-SFA. This facilitated bridging the gap between *in vivo* and *in vitro* responses as well as between static and dynamic inputs, mitigating these concerns. As a result, the difference in adaptation index detected between *in vitro* and *in vivo* VGS recordings is probably attributed to the disparity in inputs (step vs synaptic), with the latter being most likely influenced by the mixed excitatory-inhibitory inputs (including Inhibition of Inhibition, IoI) the population of NS model neurons receive. Although synaptic bombardment can lead to a high-conductance states *in vivo*, the addition of such synaptic bombardment to the population of NS model neurons was insufficient to capture the E-SFA of NS units *in vivo* (**Fig. S3**). These results suggest that while high conductance states due to *in vivo* synaptic bombardment and perhaps neuromodulation confer different integrative properties than those *in vitro*, enabling neurons to track their synaptic inputs with high temporal precision (Destexhe et al., 2001; Destexhe et al., 2003; Paré et al., 1998), such high conductance states do not interact with the mechanisms generating adaptation *in vivo*.

### Role of Local Inhibition of Inhibition

In mouse PFC, it was previously shown that parvalbumin positive (PV) neurons inhibit other PV neurons (Kvitsiani et al., 2013), somatostatin (SOM) neurons inhibit other SOM neurons (Kvitsiani et al., 2013), and vasoactive intestinal peptide (VIP) neurons inhibit PV and SOM neurons (Pi et al., 2013). However, species-specific differences in prefrontal cortex interneurons are known to exist between macaques and rodents, notably with macaque PV interneurons being more excitable than those in rat PFC (Povysheva et al., 2008). Despite these inter-species differences, fast spiking (FS) interneurons in both rat and macaque PFC have similar functional properties that enable feedforward inhibition, suggesting evolutionary invariance in circuit roles of FS interneurons across species (Povysheva et al., 2006). Here, we incorporated a common inhibitory circuit motif (IoI) present in the prefrontal cortex (Kvitsiani et al., 2013; Pi et al., 2013; Povysheva et al., 2006) into our hybrid data-driven modeling approach as a mechanism to bridge the gap between *in vitro* and *in vivo* recordings. Interestingly, macaque prefrontal cortex has an increased proportion of Calretinin (CR) positive interneurons relative to PV and Calbindin (CB) positive cells (Gabbott & Bacon, 1996; Ma et al., 2013; Medalla et al., 2023; Melchitzky et al., 2005; Torres-Gomez et al., 2020). According to their role in the circuit dynamics, CR and CB cells are considered functional homologues to VIP and SOM neurons in the mouse (Medalla et al., 2023). It is possible that CR neurons provide the IoI inputs that stabilize the response profiles of the NS neurons. While other inhibitory circuit motifs exist and cannot be ruled out from having a role in shaping *in vivo* NS neuronal response during the VGS task, the presence of such motifs further emphasizes the crucial role of IoI. Overall, the interactions between intrinsic neuronal properties, such as I-SFA, and circuit connectivity are key in the understanding of *in vivo* responses of the LPFC during a VGS task.

### Bridging the Gap between I-SFA in vitro and E-SFA in vivo

Adaptation has been documented in the *in vivo* responses of visual neurons across cortical areas of non-human primates (Muller et al., 1999; Priebe et al., 2002) and has been modeled as an intrinsic property of the neurons (Liu & Wang, 2001; Schwabe et al., 2001). Indeed, intrinsic SFA has been hypothesized to play a fundamental role in neural computations underlying high level functions (Salaj et al., 2021). The integration of I-SFA into models consisting of populations of integrate and fire neurons aided in replicating the neuronal activity recorded *in vivo (Buonocore et al., 2016; Jolivet et al., 2005; Liu & Wang, 2001)*, further supporting this notion. However, to our knowledge, detailed biophysical modelling of intrinsic SFA to understand its contribution and role in the activity of LPFC recorded *in vivo* is lacking. Furthermore, biophysical and data-driven modelling approaches of macaque neuronal activity tend to focus on understanding intrinsic neuronal properties *in vitro* or replicating *in vivo* data but not linking the two modalities. For example, the consequences of age-related morphological changes in layer 3 LPFC neurons of rhesus monkeys has been assessed using multicompartmental Hodgkin-Huxley type models and can reproduce experimentally measured increases in firing rates (Coskren et al., 2015). However, the linking of these changes in the intrinsic properties of neurons to the function of the LPFC during animal behavior was not performed using these models. Similarly, Dura-Bernal et al. (2023) have developed a data-driven model of the macaque auditory thalamocortical circuit that can replicate *in vivo* LFP and current source density data, but this model was not used in the investigation of how intrinsic neuronal properties translates into *in vivo* activity, although this potential was raised as promising future direction (Dura-Bernal et al., 2023). Experimentally, Tan et al. (2014) have performed whole-cell patch clamp recordings in V1 of behaving macaque monkeys, highlighting the exciting possibility of measuring intrinsic neuronal properties in macaque LPFC *in vivo* in the future.

### Role of SFA in PFC Function

Inhibitory cortical layer 2/3 neurons receive excitatory input in a layer-specific manner in the mouse somatosensory cortex (Xu & Callaway, 2009). If inhibitory cortical neurons solely contribute to the local circuit inhibition (both feedforward and feedback) by altering the sign of signals (i.e. transitioning from excitation to inhibition), they would be expected to convey the dynamics of inputs with high fidelity. As a result, E-SFA seen in NS neurons would be the consequence of the adaptation in the excitatory input. However, the interaction between excitatory input and intrinsic SFA demonstrated here directly contradicts this exclusively sign-change role and demonstrates that the intrinsic properties of NS neurons contribute to their activity *in vivo.* A recent study has demonstrated that LPFC neurons encode memory episodes via neural activation sequences (NAS) (Busch et al., 2024). NAS are sequences of transient activations of pyramidal cells, that last less than 500ms. Interestingly, suppression of NS cell firing using the NMDA antagonist Ketamine interfered with NAS suggesting that inhibition plays a central role in this circuit mechanism (Roussy, Luna, et al., 2021). One possible role of E-SFA is to facilitate the emergence of NAS by facilitating transient of pyramidal cells encoding the features of a memory within the LPFC circuitry. In a more general framework, E-SFA would act as a high pass filtering mechanism that allows the LPFC circuitry encoding perceptual or mnemonic events with high temporal resolution.

### Narrow Spiking Neurons are Mostly Fast Spiking Interneurons

The firing rate of NS neurons is higher than that of BS neurons *in vitro* (**Figure 4B** and **F**), consistent with being fast-spiking interneurons. Additionally, the majority (20/33) of NS neurons recorded *in vitro* are classified as fast-spiking based on their I-SFA and the Petilla classification (Ascoli et al., 2008). Additionally, the *in vitro* NS neuron adaptation index values (0.838 (IQR 0.620-0.935)) are in line with those of linear arbor cells (0.89±0.2) (Zaitsev et al., 2009), and chandelier cells (0.83±0.2 and 0.86±0.19) (Povysheva et al., 2013; Zaitsev et al., 2009) fast spiking parvalbumin positive interneurons in macaque LPFC. However, other quantifications of I-SFA in fast spiking primate LPFC interneurons have found larger mean adaptation indices than reported here (González-Burgos et al., 2005; Povysheva et al., 2006; Povysheva et al., 2008). As a result, the NS neuron populations *in vitro* and likely *in vivo* probably contain a heterogeneous collection of interneurons, despite classification based on spike width. If the dependence of *in vivo* E-SFA on IoI and its interaction with I-SFA occurs with different prevalence or impact across different interneuron types remains an area for future research. Despite these differences in magnitude of I-SFA and potential heterogeneity of the NS populations, I-SFA is present in NS neurons presented here and in FS interneurons (Ascoli et al., 2008; González-Burgos et al., 2005; Povysheva et al., 2006; Povysheva et al., 2013; Povysheva et al., 2008; Zaitsev et al., 2009) and as such the timescale of adaptation can be quantified regardless of the magnitude of adaptation.

### SFA as a Target for Neuromodulation

Increased magnitude of E-SFA *in vivo* compared to I-SFA *in vitro* has been suggested to result from differences in extracellular ionic environment and/or the presence of neuromodulators (Fernandez et al., 2018). Many neuromodulators can affect I-SFA mechanisms such as calcium sensitive small conductance potassium (SK) channels (Faber et al., 2008; Kramar et al., 2004; Lin et al., 2008; Maingret et al., 2008; Ren et al., 2006) and slow non-inactivating potassium M-current (Brown, 1988; Cooper & Jan, 2003). As such, modulation of intrinsic SFA through modulation of calcium activated potassium channels and/or M-currents could serve as a mechanism through which neuromodulators exert their effects on PFC neuronal firing, circuitry and function. However, the effects of neuromodulators, to our knowledge, generally involve modulation of I-SFA current amplitudes (attenuation/enhancement) and not of the timescale of the I-SFA. As such we view it as unlikely that neuromodulators could replace the role of IoI *in vivo* described here.

### Limitations

Although the hybrid modelling approach presented here was driven by *in vivo* BS activity and examined possible local circuitry, the extensive long-range connectivity of the LPFC (Haber et al., 2022; Saleem et al., 2014; Yeterian et al., 2012) was explicitly neglected here. However, we employed the data-driven BS neural responses as excitatory input into the NS model neurons, allowing us to implicitly account for the effects of some of these inter-area connections onto BS neurons should they be present.

### Conclusion

Our results reveal that macaque LPFC BS and NS neurons undergo adaptation during *in vivo* and *in vitro* experiments, despite receiving visual stimulus driven input from intact circuitry *in vivo* and receiving step-current inputs without synaptic connectivity *in vitro.* A data-driven hybrid modelling approach was then used to explore how the intrinsic SFA contributes to the *in vivo* adaptation. A local inhibitory component resulting from circuit dynamics (IoI) interacting with intrinsic SFA properties in a timescale specific manner successfully reproduced the *in vivo* adaptation. Thus, adaptation in LPFC neurons during behavior results from the interaction between intrinsic neuronal properties and inhibitory circuits dynamics. The latter highlights the combined role of single neuron and network computations on generating the patterns of neural activity underlying behavior.

## Acknowledgements

This work was supported by the Natural Sciences and Engineering Research Council of Canada (NSERC) Discovery Grant to A.K., Canadian Institutes of Health Research (CIHR), NSERC, and BrainsCAN grants to J.C.M-T., a Neuronex (ref. FL6GV84CKN57) grant to S.T., J.F.S., G.G.B., A.N., S.T., S.E., W.I., and J.C.M-T. and a NSERC Postgraduate Scholarship-Doctoral Fellowship to N.A.K..

## Author Contributions

Conceptualization: N.A.K., B.W.C., A.K., and J.C.M-T.; Data curation: N.A.K., B.W.C. and M.F.; Formal Analysis: N.A.K.; Investigation: N.A.K., B.W.C., M.F., R.A.G., M.S.J-S., M.A., J.K.S., S.M., M.R., R.L.A., S.A.M., B.M., S.V., H.I., K.S.P., and W.J.A.; Methodology: N.A.K., B.W.C., A.K., and J.C.M-T.; Software: N.A.K.; Visualization: N.A.K.; Writing - original draft: N.A.K. and B.W.C.; Writing - review & editing: N.A.K., B.W.C., M.F., A.P., S.T., J.F.S., G.G.B, A.N., S.T., S.E., W.I., A.K., and J.C.M-T.; Funding acquisition: A.K., S.T., J.F.S., G.G.B., A.N., S.T., S.E., W.I., and J.C.M-T.; Resources: A.P., S.T., J.F.S., G.G.B., A.N., S.T., S.E., W.I., A.K., and J.C.M-T.; Supervision: A.P., S.T, J.F.S., G.G.B, A.N., S.T, S.E., W.I., A.K., and J.C.M-T.;

