## Supplemental Material for "Spike frequency adaptation in primate lateral prefrontal cortex neurons results from interplay between intrinsic properties and circuit dynamics"

**Supplementary Figures**

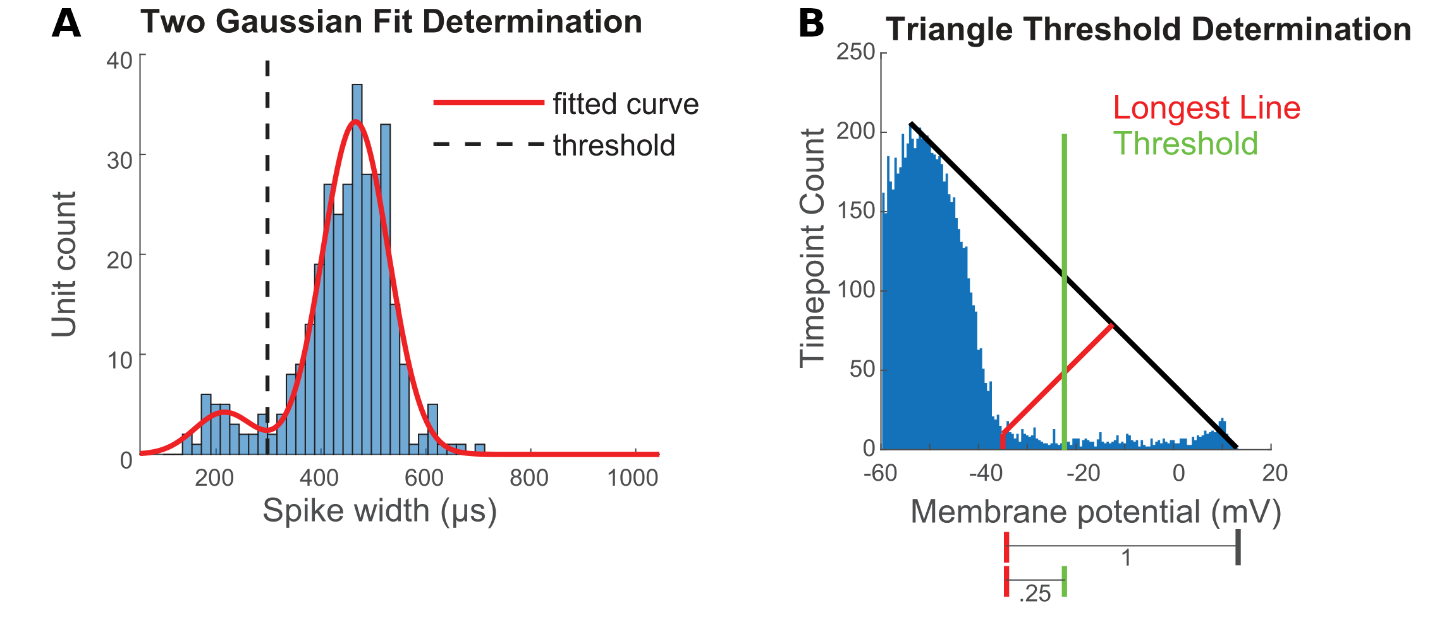

**Figure S1**. (A) The sum of two Gaussians (red line) fitted to the distribution of *in vivo* spike widths (blue histogram); the local minima between the two Gaussians (identified by the dashed line) is utilized to differentiate between NS and BS cells located to the left and right of the dashed line, respectively. (B) The Zack et al. (1977) triangle method utilized to define the threshold for spike detection. This method employs binning to initially create a distribution of the membrane voltage. Subsequently, it delineates the threshold by first establishing a line connecting the highest bin count to the bin with the highest value (black line). Then it identifies the longest perpendicular line from the black line to each bin (red line). The bin associated with the longest perpendicular line corresponds to the critical value. Finally, the threshold voltage is determined as one-quarter of the distance between the critical value and the maximum voltage value, as indicated by horizontal lines below the graph.

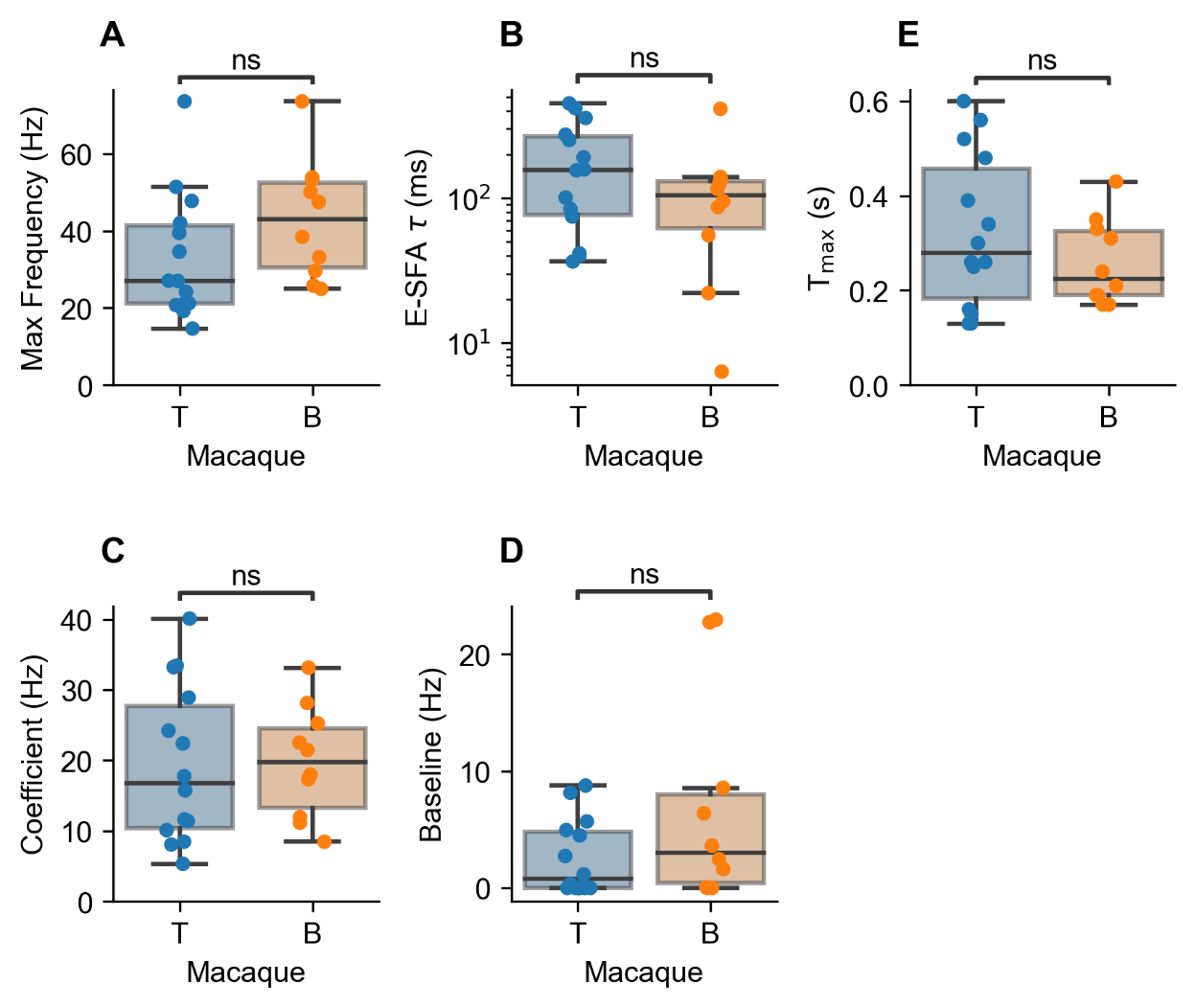

**Figure S2**. No differences in various features of adaptation and latencies of broad spiking units between animals. The distributions of maximal (A) firing frequencies, (B) embedded SFA (E-SFA) decay time constants, (C) exponential decay coefficients, (D) exponential decay baselines, and (E) latencies to maximal PSTH are not significantly different. The E-SFA decay time constants, coefficients and baselines were computed using Eq. (2). ns p≥0.05, * p < 0.05, **p<0.01, *** p < 0.001

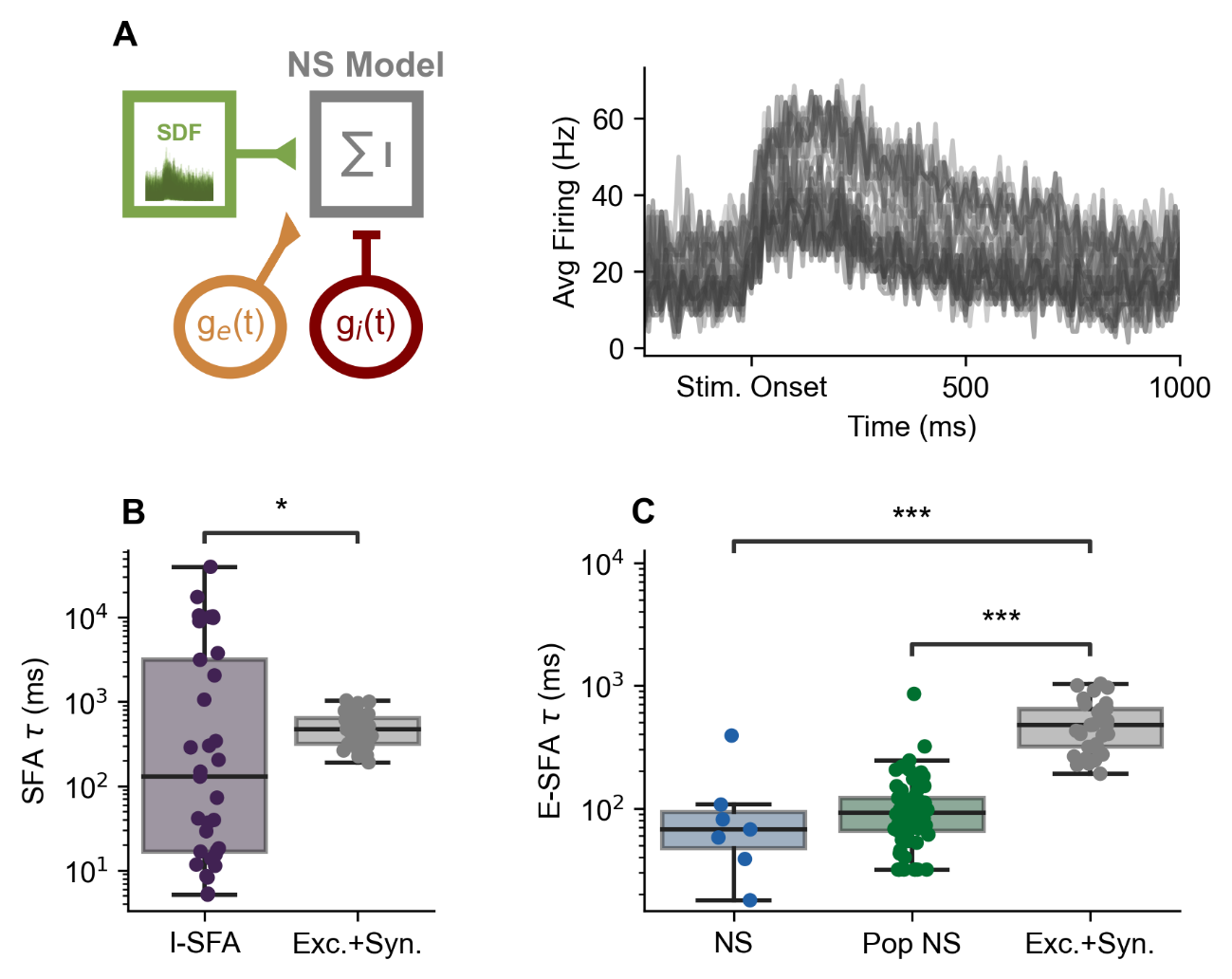

**Figure S3.** Responses of a population of NS model neurons to a combination of BS input and synaptic bombardment. (A) Left: schematic depiction of a hybrid circuit model with Ornstein Uhlenbeck excitatory (g_E_(t)) and inhibitory (g_I_(t)) synaptic input added as synaptic bombardment. Excitatory input from *in vivo* BS units (green) and the synaptic bombardment were injected as inputs to a population of NS model neurons (grey). Right: the computed PSTHs generated by these NS model neurons. (B) Comparison of model decay time constants (computed by fitting Eq. (2)) in response to the step-current stimulus (3µA/cm^2^; I-SFA) against those produced by the model in (A) with synaptic bombardment (Exc.+Syn.; Mann-Whitney, U= 387.0, p= 0.044). (C) Comparison of the *in vivo* NS (NS) and population level NS (Pop NS) time constants (computed by fitting Eq. (2)) against those produced by the model in (A) when subjected to synaptic bombardment (Exc.+Syn.; Kruskal Wallis H-test, H=62.23, p= 2.176e-14). Dunn’s post hoc test p-values are reported after a Kruskal-Wallis tests. ns p≥0.05, * p < 0.05, **p<0.01, *** p < 0.001

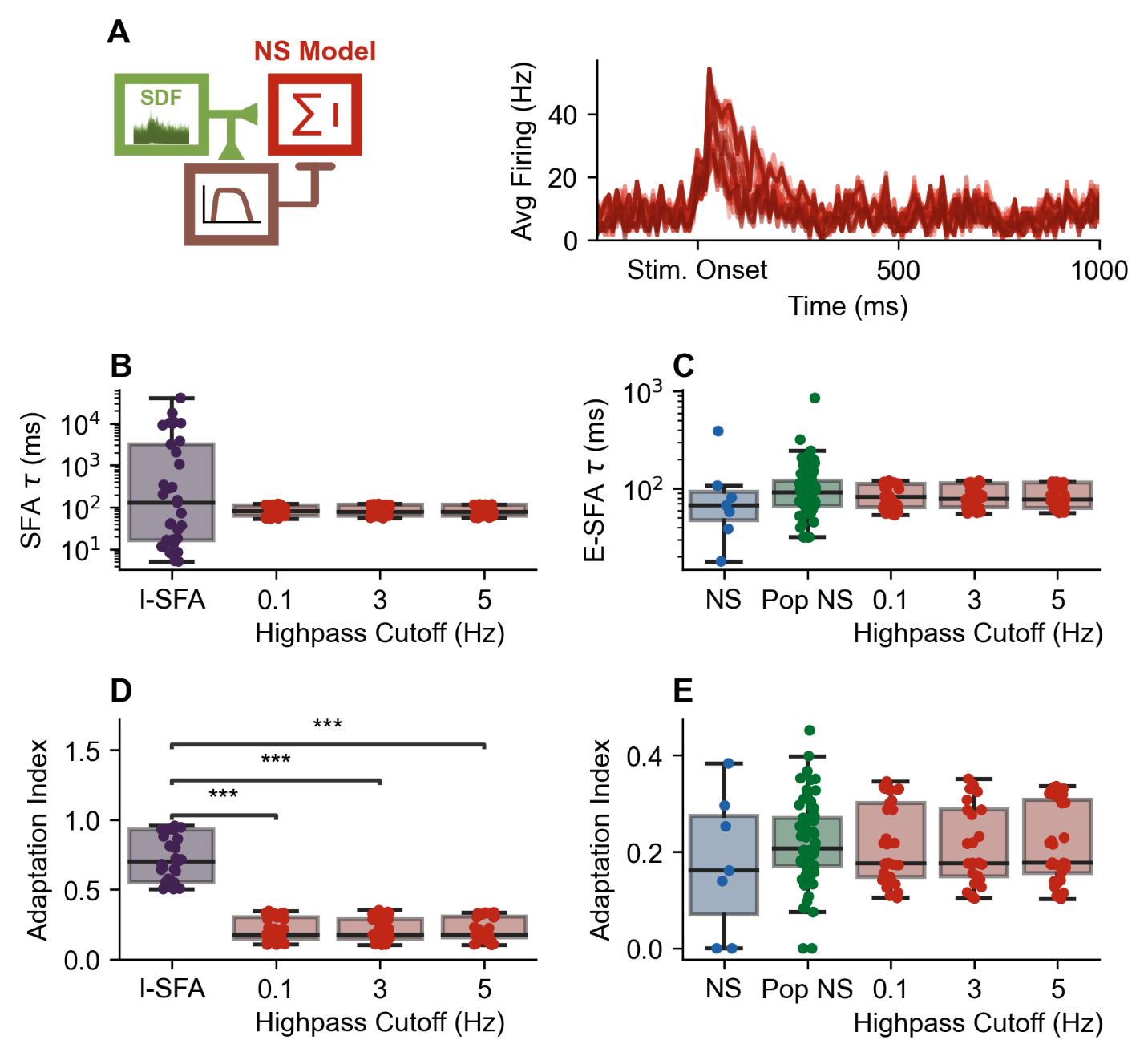
**Figure S4.** Responses of a population of NS model neurons to a combination of BS input and feedforward inhibition with band-pass properties. (A) Left: schematic depiction of a hybrid circuit model with high-pass SFA added to produce a band-pass feedforward inhibition (brown, 0.1 Hz high-pass cutoff). Excitatory input from *in vivo* BS units (spike density function (SDF); green) and feedforward inhibition were injected as inputs to a population of NS model neurons (red). Right: the computed PSTHs generated by these NS model neurons. (B) Comparison of model decay time constants (computed by fitting Eq. (2)) in response to the step-current stimulus (3µA/cm^2^; I-SFA) against those produced by the model in (A) when inhibition was subjected to a high-pass filter with cutoff: 0.1, 3 and 5 Hz (Kruskal Wallis H-test, H=0.1265, p=0.988. (C) Comparison of the *in vivo* NS (NS) and population level NS (Pop NS) time constants (computed by fitting Eq. (2)) against those produced by the model in (A) when inhibition was subjected to a high-pass filter with cutoff: 0.1, 3 and 5 Hz (Kruskal Wallis H-test, H=4.692, p=0.320). (D) Comparison of model adaptation indices in response to the step input current (3µA/cm^2^; I-SFA) against those produced by the model in (A) when inhibition was subjected to a high-pass filter with cutoff: 0.1, 3 and 5 Hz (Kruskal Wallis H-test, H=73.725, p=6.80*10^-16^). (E) Comparison of the *in vivo* NS (NS) and population level NS (Pop NS) adaptation indices against those produced by the model in (A) when inhibition was subjected to a high-pass filter with cutoff: 0.1, 3 and 5 Hz (Kruskal Wallis H-test, H=7.131, p=0.129). Dunn’s post hoc test p-values are reported after a Kruskal-Wallis tests. ns p≥0.05, * p < 0.05, **p<0.01, *** p < 0.001

**Supplementary Tables**

| **Patch NS** $\boldsymbol{\tau}$ **(ms)** | **Fit model** $\boldsymbol{\tau}_{\boldsymbol{max}}$ **(ms)** |
| --- | --- |
| 15.234 | 235.685 |
| 2053.981 | 30140.436 |
| 13.737 | 212.422 |
| 10138.804 | 90765.190 |
| 8.186 | 124.664 |
| 5.310 | 75.075 |
| 39.621 | 590.317 |
| 3849.211 | 51295.463 |
| 72.778 | 1073.243 |
| 287.154 | 4253.096 |
| 8.544 | 129.413 |
| 18979.322 | 112468.184 |
| 17.099 | 262.330 |
| 60507.223 | 131381.758 |
| 9497.588 | 87770.863 |
| 204.434 | 3012.464 |
| 11.687 | 180.995 |
| 5.152 | 73.141 |
| 18.268 | 279.552 |
| 1057.790 | 15984.377 |
| 36.867 | 550.756 |
| 29.138 | 439.331 |
| 3166.415 | 44140.621 |
| 148.020 | 2176.595 |
| 342.193 | 5087.949 |
| 11310.757 | 93682.419 |
| 11.347 | 175.392 |
| 9853.853 | 91657.052 |
| 301.117 | 4465.878 |
| 10670.522 | 92916.097 |
| 16.715 | 257.180 |
| 129.364 | 1901.733 |
| 41.388 | 616.458 |

**Table S1.** Adaptation current $\tau_{max}$ parameter fits to reproduce *in vitro* patch clamp SFA time constants.
